# Trait distribution in Geometrid moth assemblages does not vary along a western Himalayan elevational gradient

**DOI:** 10.1101/2024.04.01.587530

**Authors:** Pritha Dey

## Abstract

Elevation gradients pose challenges for flying insects due to temperature and air density shifts, impacting their thermoregulation, flight abilities and subsequently species distribution patterns. Geometrid moths are a model group for studying these traits among insects. This study examines how trait-patterns and niche differentiation of Geometrid moth assemblages change with elevation. Investigating 697 specimens of 120 Geometrid moth species, I assessed species diversity, turnover, and various traits related to body size and flight capabilities along a 1500m elevation span in the western Himalayas. Results indicate a decline in species diversity along the elevational gradients, accompanied by consistent turnover from lower to higher elevations. Despite this pattern, elevation did not significantly influence morphological traits related to thermal sensitivity (body size) and flight capability (wing loading and manoeuvrability) at the assemblage level in Geometrid moths. Furthermore, there was no significant niche differentiation among species across elevations. This supports the argument that species with similar ecological roles persist through turnover, maintaining consistency in trait values across assemblages. These findings from the paleotropics align with patterns observed in Geometrid moths across neotropical and temperate regions, highlighting the importance of trait-based approaches in understanding species responses to environmental changes.

**Summary Statement:** This study examines how Geometrid moth assemblages in the western Himalayas respond to elevational gradients, revealing consistent turnover, stable traits, and no significant niche differentiation, emphasizing trait-based species adaptation

## Introduction

Mountain ecosystems are increasingly threatened by climate change, posing a risk to cold-adapted and endemic species that depend on these regions (Laiolo *et al.,* 2018; Trew & Maclean, 2021). These unique ecosystems have drastic environmental changes over small spatial scales (Rogora *et al.,* 2018) affecting species distributions across gradients. To survive these conditions, species need to adapt physiologically, which is often reflected their morphological traits (Ryding *et al*., 2021; Warrington & Waterman, 2023). Morphological traits are, therefore, crucial for our understanding of species assembly (Hoiss *et al*., 2012; Chichorro *et al*., 2022). The importance of morphological traits is enhanced while studying assemblages of species rich clades such as insects, where species identification is challenging (Yates *et al.,* 2014)

Insects adapt to abiotic factors such as elevation, temperature, and air density indirectly through interactions with hostplants, competitor species, parasitoids and predators (Hodkinson, 2005), or directly through traits like body size, wing size and colouration (Xing *et al*., 2018; Bladon *et al*., 2020; Clusella-Trullas & Nielsen, 2020). Body size (often evaluated as wingspan), in particular influences thermoregulation, and is a key focus in studies on patterns of traits in insects across elevation and latitudes (reviewed in Chown & Gaston, 2010). For example, bees decrease in size with elevation (Osorio-Canadas *et al*., 2022), but increase across latitudes (Gérard *et al.,* 2018). Dragonflies and damselflies are larger in temperate regions than in the tropics (Svensson *et al*., 2023). Noctuid moths show larger body sizes at higher elevations in temperate regions (Heidrich *et al*., 2021). However, there is significant variability among taxa. For instance, Geometrid moths, show diverse patterns: some studies indicate a positive relationship between body size and elevation (Beck *et al.,* 2016; Brehm *et al.,* 2018), while others reveal no change (Brehm & Fiedler, 2004). Additionally, body size directly impacts flight-related traits (Grula *et al.,* 2021), with larger insects having higher wing loading and lower manoevrability (Shyy *et al.,* 2016). At higher elevations, lower temperatures favour species with smaller body and reduced wing loading (therefore proportionally long wings) (Dudley, 2002) which is advantageous in low air density (Dillon & Dudley, 2004; Dillon *et al*., 2006; Grula *et al*., 2021).

Morphological traits are critical in defining a species’ niche in an assemblage (Violle & Jiang, 2009) and inferences can be made regarding community assembly by measuring changes in the niche space and trait overlap across ecological gradients. At lower elevations, warm temperature and abundant resources reduce constraints, leading to high diversity and niche specialization with lower trait overlap (Read *et al*., 2018), as seen in Orthopteran (König *et al.,* 2024) and bee communities (Hoiss *et al.,* 2012). Conversely, higher elevations impose stricter environmental constraints, resulting in lower diversity and higher trait overlap, consistent with the ‘niche packing’ model (Macarthur,1965; Pigot *et al.,* 2016; Pellissier *et al.,* 2018). The degree of trait overlap is shaped by niche differentiation and the displacement of functional trait values over evolutionary time (Macarthur & Levins, 1967; Ricklefs & O’Rourke, 1975). However, distinguishing traits affected by biotic versus abiotic environmental filters, in trait-based approaches, can be challenging (Violle & Jiang, 2009). Analyzing the traits weighed by the relative abundance of the species best reflect the adaptation of traits to the environment (Garnier *et al.,* 2004; Shipley *et al.,* 2006). Community Assembly Theory (HilleRisLambers *et al.,* 2012; Kraft *et al.,* 2015) predicts that assemblages that are shaped by interspecific competition show overdispersed trait spaces whereas those shaped by environmental adaptations are underdispersed, given the relatedness of the species in the assemblage (Weiher & Keddy, 1995)

In this context, I Firstly wanted to understand 1) how does species diversity and turnover change across the elevation? If there is a change in species diversity across elevation, 2) do body size and flight related traits show a predictable relationship with elevation?, and 3) do trait dispersion and the overall trait overlap (niche differentiation) at assemblage level vary with changing species diversity along the elevation? I predict species diversity to decrease with high turnover with increasing elevation. I hypothesize that body size and flight traits are expected to change with elevation to achieve optimal thermoregulation and flight, leading to reduced body size, reduced wing loading and higher manoevrability. Following species assembly rules, higher elevations are expected to show trait under dispersion, higher trait overlap, with less niche differentiation due to decreased species diversity. While these patterns have been observed in bird and bat communities (Schumm *et al.,* 2020; Chakravarty *et al.,* 2023), in the tropical Himalayan mountain system, it remains unexplored if similar patterns apply to insects as well.

I studied Geometrid moth assemblages across a 1500m elevation gradient on the fringes of Kedarnath Wildlife Sanctuary. Despite the extensive use of Geometrid moths in ecological studies (Brehm & Fiedler, 2004; Davis *et al.,* 2013; Beck *et al.,* 2016; Holm *et al.,* 2019a, 2019b; Choi *et al.,* 2022; Seifert *et al*., 2022; Foerster *et al.,* 2024) so far no study has considered body size and flight-related traits together to understand the effect of elevation on the overall trait space and the niche differentiation. This trait-based approach aims to fill this gap, enhancing our ability to address ecological questions, particularly given the unclear phylogenetic positions of Geometrid moths from this region in global studies.

Furthermore, research on macroecological patterns within paleotropical montane systems is limited. While previous studies from the western Himalayan region, Sanyal et al. (2017) and Dey (2019) have investigated species diversity patterns across elevational gradients, they did not consider trait-based patterns. Trait-based research on moths from this region focused on less diverse groups like Hawkmoths, highlighting significant trait variations and internal filters influencing community assembly (Mungee & Athreya 2021). This study specifically focused on, on understanding how elevation and related abiotic factors influence trait-level patterns among highly diverse Geometrid moths.

## Results

### Species diversity

Out of 119, there were 56 species level identifications, 56 morphospecies identified only till genus, one was identified till subfamily, three morphospecies were identified till tribe and three morphospecies remain unidentified (Table S3). Figure 1b illustrates a decrease in species diversity at higher elevations. Rarefaction curves facilitate the comparison of diversity across various communities or habitats. By scrutinizing the shapes and positions of these curves (Figure 1a and 1b), the disparities in species diversity, sampling completeness, and overall species diversity among sites are clear. A steep slope indicates rapid accumulation of new species with additional sampling effort, suggesting high species richness and/or incomplete sampling. In contrast, a shallow slope indicates that most species have already been sampled, and further sampling is unlikely to yield many new species.

**Figure 1:**
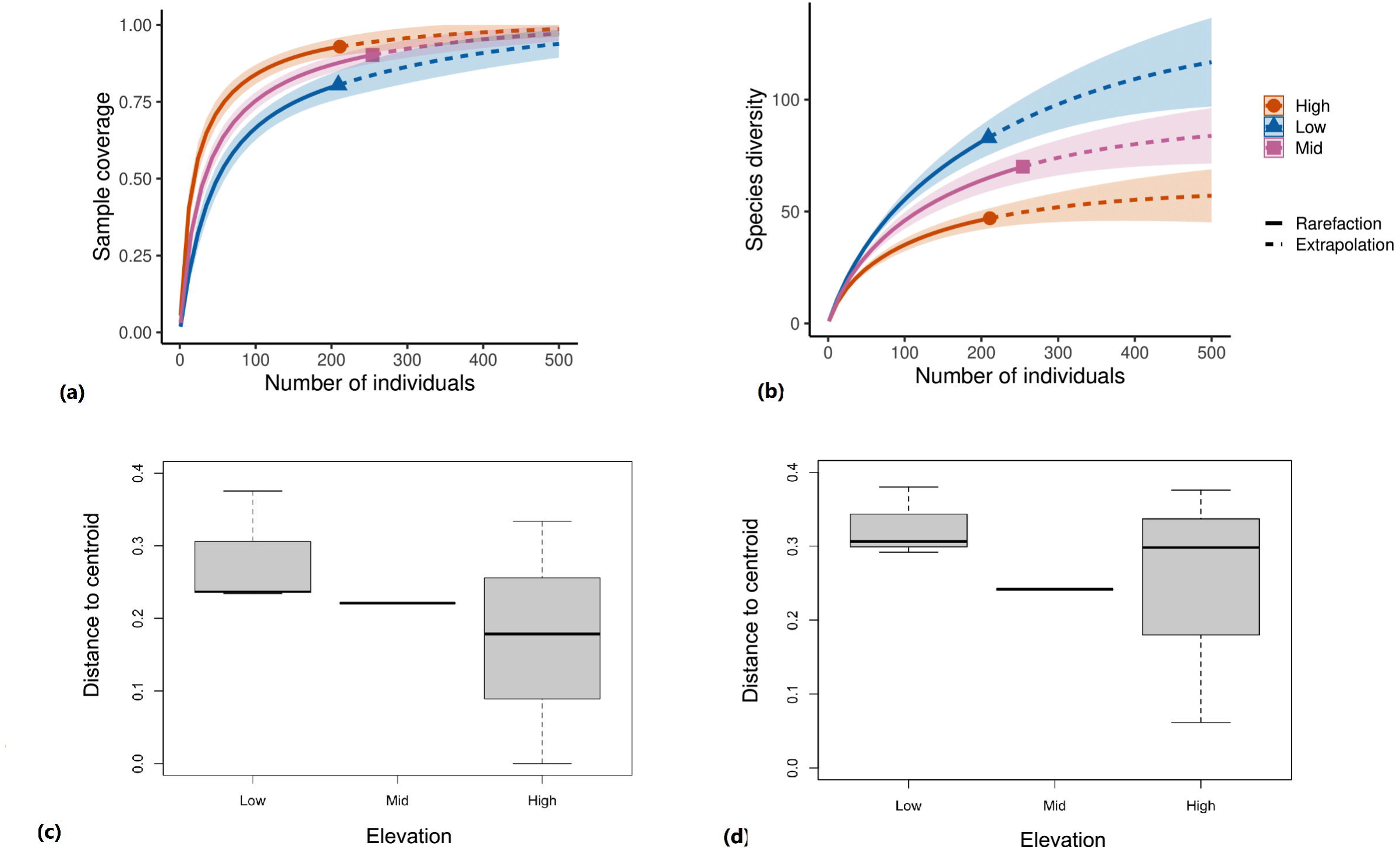
Rarefied curves for a) Sample coverage and b) Species diversity patterns extrapolated across the number of individuals; c) overall beta diversity measured as Sorensen pairwise dissimilarity and d) the spatial turnover of the species measured as Simpson pairwise dissimilarity across the low, mid and high elevation assemblages

For beta diversity, the resulting box-plot shows the distribution of βnested and βturnover values across the different pairs of sites or groups in your data (Fig 1c and Fig 1d). This visualization helps to understand the relative contributions of nestedness and turnover to the overall beta diversity pattern. The Euclidean distance to centroid for Sørensen pairwise dissimilarity among the elevation categories were as follows: Low-Mid (0.314); Low-High (0.489) and Mid-High (0.34); similarly, the Simpson dissimilarity index (spatial turnover) was: Low-Mid (0.05); Low-High (0.14) and Mid-High (0.12) (Fig 1 c and d). Though there is indication of dissimilarity in species composition and turnover of species between the low and high elevation sites, the Euclidean distances are not significantly different from each other (Sørensen pairwise dissimilarity: ANOVA, df=2, p>0.05; Spatial turnover: ANOVA, df=2, p>0.05)

### Variation in the individual trait values, trait dispersion and trait space overlap (niche differentiation)

There was no significant effect of elevation on the community-weighted means of the individual trait values (Fig.2) (model summary provided in Table S2). The pairwise comparisons of the raw means of the individual traits across the elevation classes also did not show any significant difference (details in Figure S3). There is high variability and a weak pattern in the CWM across elevation, which makes it evident that the patterns are sensitive to the abundance of species. The species of same trait values are abundant across the elevation. The density plots of individual traits would make it clearer, where kernel density estimate is used to show the distribution of each of the traits, indicating the mean values (Figure S4).

**Figure 2:**
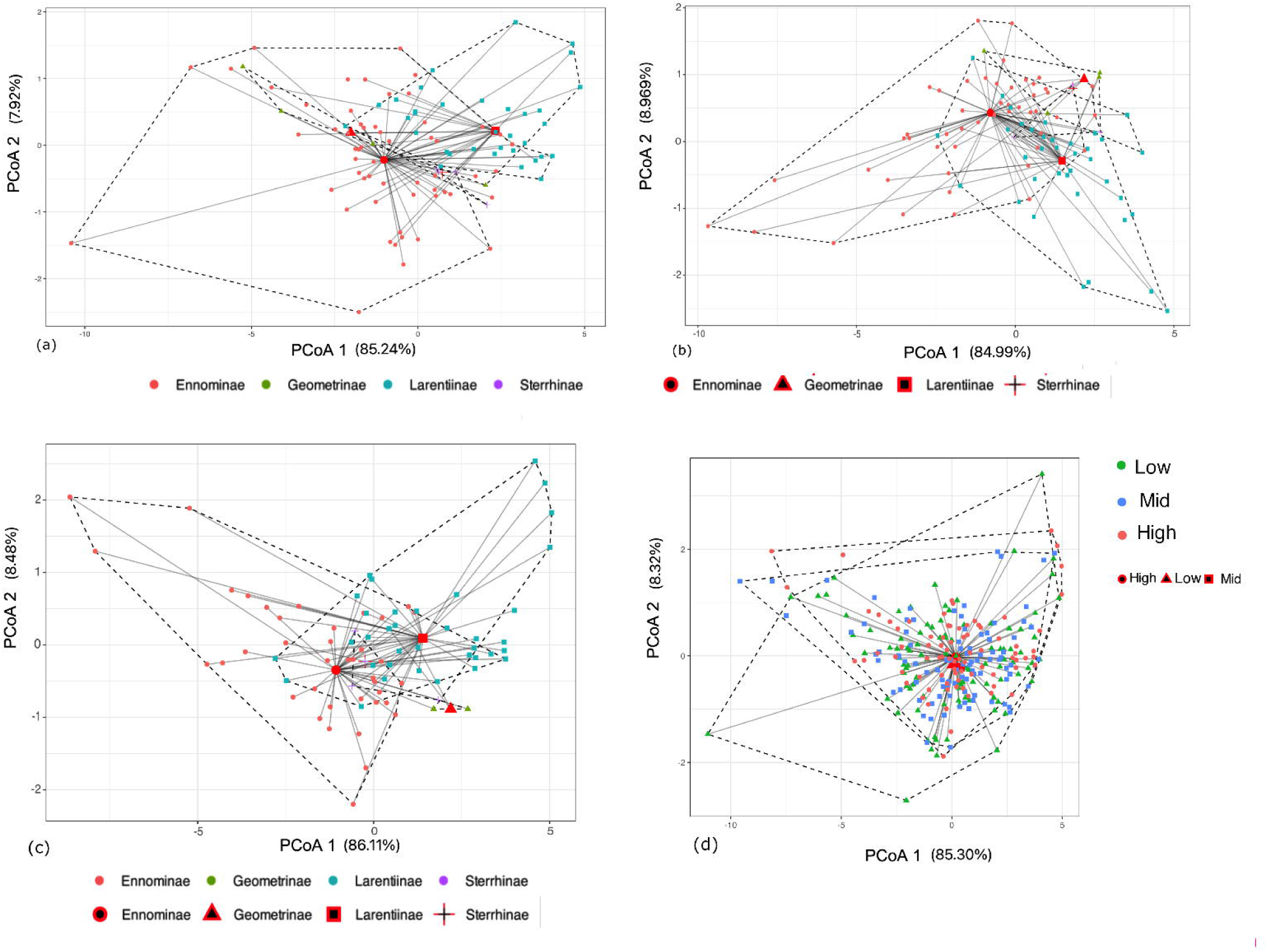
The figure shows the community weighted means (mean ± SE) of each of the traits across the low, mid and high-elevation assemblages.

Trait dispersion did not differ significantly among the subfamilies in the low (df=3, p=0.149), mid (df=3, p=0.252), and in high elevation (df=3, p=0.2). Similarly, no significant differences were found in trait dispersion among assemblages across the elevation (df=2,p=0.3) (Fig.3).

**Figure 3:**
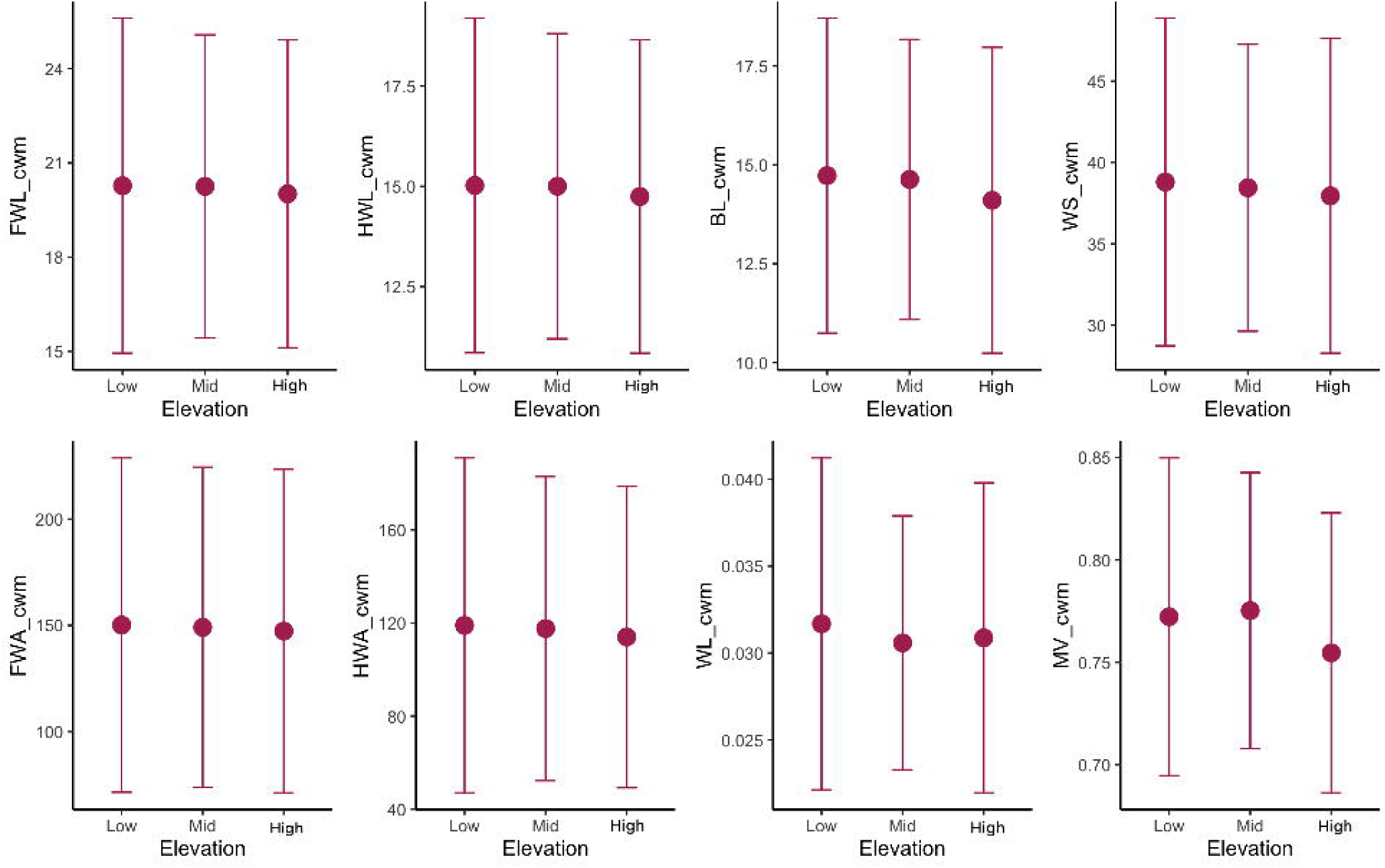
The figure shows the comparison of subfamily-level trait dispersion in a) Low, b) Mid and c) High elevation classes; and overall comparison of trait dispersion at the d) assemblage level. The red-black shapes denote the centroid for the respective level of comparison.

There is also a high level of niche overlap among the species of the Geometrid moth communities across the elevation, as shown by high (0.84) O-statistic index of the multivariate trait space (Fig. 4). This indicates that for each trait the observed niche overlap did not differ from that which would be expected due to chance.

**Figure 4:**
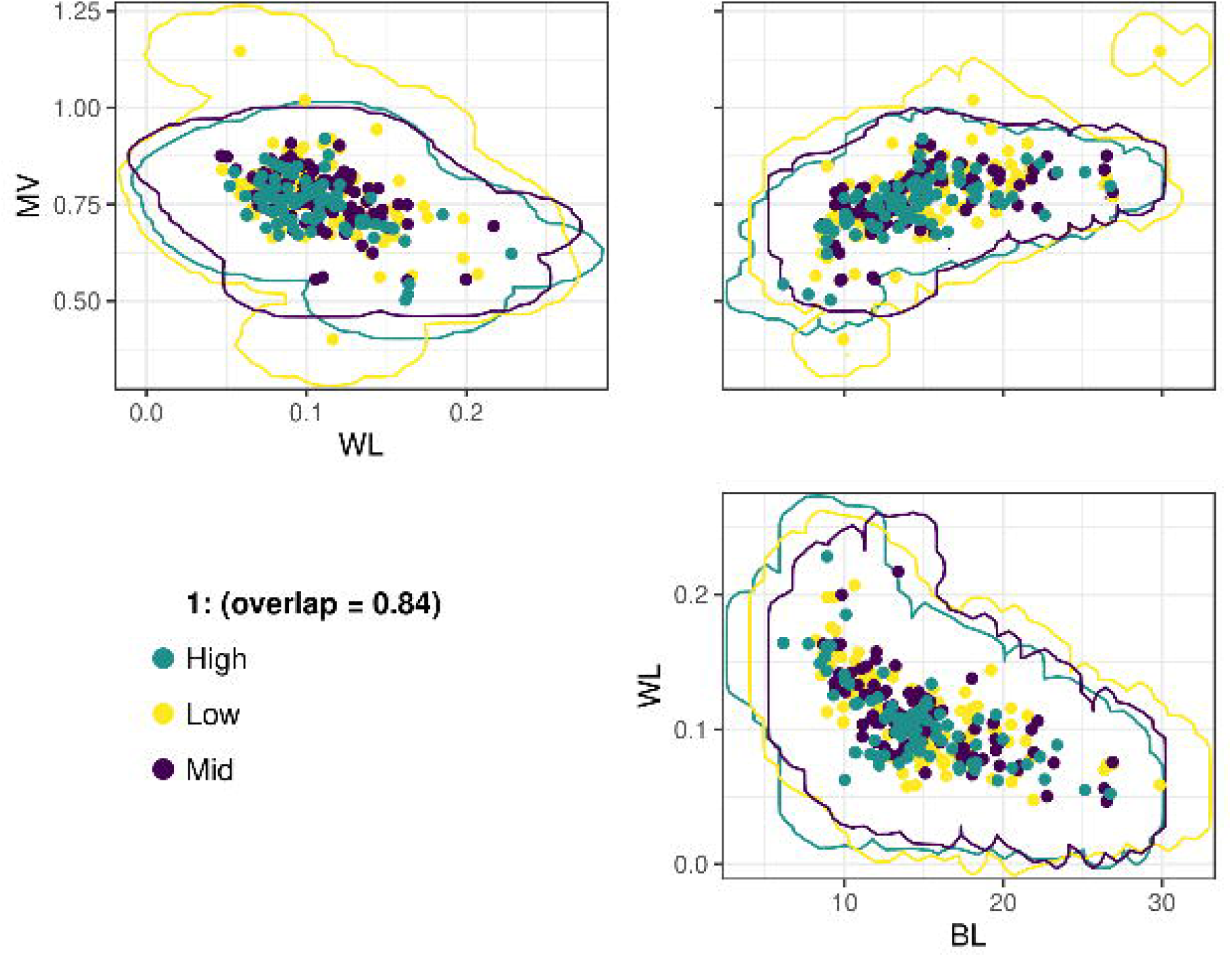
The figure shows the degree of overlap (O-statistic) of the multivariate trait-space, based on body length (BL), maneuverability (MV), and wing loading (WL) of the Geometrid moth assemblages in low, mid, and high elevations

## Discussion

The findings offer important insights into the dynamics of Geometrid moth assemblages along a 1500m gradient in the western Himalaya. The results show a decrease in species diversity with increasing elevation, along with consistent species turnover from the low to the high elevations. Despite this, elevation does not significantly impact morphological traits related to thermal sensitivity (body size) and flight capability (wing loading and manoevrability) at the assemblage level. This suggests that Geometrid moths maintain trait consistency across different elevations. Additionally, the study reveals no significant niche differentiation among species across the elevation.

### Species diversity across elevation

The results indicate that trait-level patterns may not directly correlate with the species diversity, and relative abundance. For instance, species from subfamily Ennominae, such as from the genera *Alcis* and *Abraxas*, were abundant at low and mid elevations, likely due to its polyphagous diet (Choi *et al.,* 2022), while species like *Dalima truncataria* were scarce. Conversely, *Psyra crypta* was more prevalent at higher elevations. The turnover from low to high elevations is influenced by the higher abundance of species from Larentiinae subfamily, particularly those specializing in herbaceous plants, reflecting similar patterns observed in other mountain gradients (Skou, 1986; Brehm & Fiedler, 2003; Brehm *et al.,* 2013). Further, local host-plant availability, diet breadth, and breeding strategies likely affects species diversity (Heidrich *et al*., 2021; Seifert *et al*., 2022). Moreover, the consistent presence and high abundance of some species across elevations might also indicate density compensation mechanisms due to competition and predator relaxation (Longino & Colwell 2011; Choi *et al*. 2022)

### Morphological trait variation across elevation

Body size significantly affects how insects respond to temperature changes, influencing their distribution and thermal tolerance (Hodkinson *et al.,* 1999; Hodkinson, 2005). However, Geometrid moths do not show significant body size changes with elevation (Bladon *et al.,* 2020; Heidrich *et al.,* 2021), likely due to their large wings and low wing loading, which allow them to fly across varying temperatures without extensive warm-up (Casey & Joos 1983; Heinrich, 2013). This wide thermal tolerance may contribute to their stable body size and flight traits across elevations. Similar adaptation patterns have been observed in other taxa, such as the African dung beetles (Gaston & Chown, 1999). The high abundance of certain common species also played a role in maintaining stability, as these species tend to have traits well-suited to the prevailing environmental conditions (Cingolani *et al.,* 2007)

The uniformity in trait dispersion among Geometrid subfamilies suggests that similar selective pressures or trait convergence are at play, despite the taxonomic diversity. Although the impact of phylogenetic constraints remains unclear due to limited taxonomic and phylogenetic knowledge of moths from this region, the minimal variation in morphological traits along elevation is likely attributed to the abundance of species with similar traits and evolutionary relatedness (Griffith *et al,* 2023.). Traits that are conserved through shared evolutionary history are apparent in many phytophagous insects. For example, closely related herbivore species often feed on the same or similar plant taxa (Nylin *et al.,* 2014; Strutzenberger *et al*., 2017) and exhibit comparable levels of dietary specialization (Slove & Janz, 2011; Hardy *et al.,* 2016) and body size (Ribeiro & Freitas, 2011). Such similarity in traits across communities, indicate a lower degree of specialization and a higher prevalence of generalist species. As a result, this increased trait similarity may reduce ecosystem resilience, affect community functioning, and impair ecosystem services (Lavorel & Garnier, 2002; Olden, 2006)

More definitive conclusions about the mechanisms driving the lack of variation in body size and flight traits in insects, and how these influence species distributions, are hindered by insufficient life-history, ecological, and abundance data collected over extended periods and with higher temporal resolution, from the Himalayan region.

### Conclusion

The study provides valuable insights into how individual traits, trait variability within the assemblages, and niche differentiation respond to changing species diversity across elevational gradients. With evidence from the Paleotropical region, the results align with the lack of consistent trends in individual traits studied in Geometrid moths and other insect taxa across diverse elevation ranges in neotropics and palearctic regions (Brehm & Fiedler, 2004; Peters *et al*., 2016; Beck *et al.,* 2016; Classen *et al.,* 2017).

No significant variation in trait-patterns with changing species diversity, challenge prevailing notions about the association between elevation and insect morphology. It highlights the possible role of functional redundancy (Pillar *et al*., 2013) or biotic homogenization (Olden, 2006)in maintaining trait distributions across varied elevations and underscores the need for a nuanced species-specific understanding of trait-environment relationships.

However, the limitations, including restricted sampling to the spring-summer season and the exclusion of lower elevation sites, suggest a need for further research. Seasonal variations and additional lower elevation sites could provide a comprehensive understanding of trait-environment dynamics. Previous studies from other western Himalayan elevational gradients, (Sanyal *et al*., 2017) and (Dey, 2019) indicate variations in species composition across seasons, highlighting the need for such an investigation. Further, integrating phylogenetic information is essential when examining species traits, their determinants, and their relationships.

Future research could explore how life history traits, such as larval host plant use (i.e., diet breadth and host-plant growth form) and seasonal life cycle (i.e., overwintering stage and caterpillar phenology); and factors like predator density are linked to adult body size and flight morphology. Such causal relationships would be crucial for developing effective conservation strategies, especially as climate change alters environmental conditions. Long-term studies investigating potential shifts in trait distributions, can provide valuable insights into how species respond to changing thermal regimes and help tailor conservation efforts to protect the unique biodiversity of the Himalayan ecosystem.

## Materials and methods

### Study Area

The study was done in 2018 around the fringes of the Kedarnath Wildlife Sanctuary (KWLS) (30°25′–30°41′N, 78° 55′–79°22′E) located in the Chamoli-Rudraprayag district in the state of Uttarakhand (Map 1). KWLS covers an area of 975 sq. km and lies in the upper catchment area of the rivers Alakananda and Mandakini, the major tributaries of the river Ganga. The elevation of the protected area ranges from 1160-7068 meters above sea level, encompassing diverse habitats from moist temperate forests, mixed oak forests to lush alpine meadows. The combination of human pressure, pristine forest areas, and a large altitudinal range make it an ideal site for biogeographic studies. The forest types are dominated by oak (*Quercus leucotrichophora, Quercus floribunda, Quercus glauca, and Quercus semecarpiflora*). The area receives annual rainfall of about 3000 mm with maximum contribution during monsoon (June to September). During the winter months (December to February) heavy snowfall takes place in the alpine and temperate regions (Thakur *et al*., 2011).

**Map 1:**
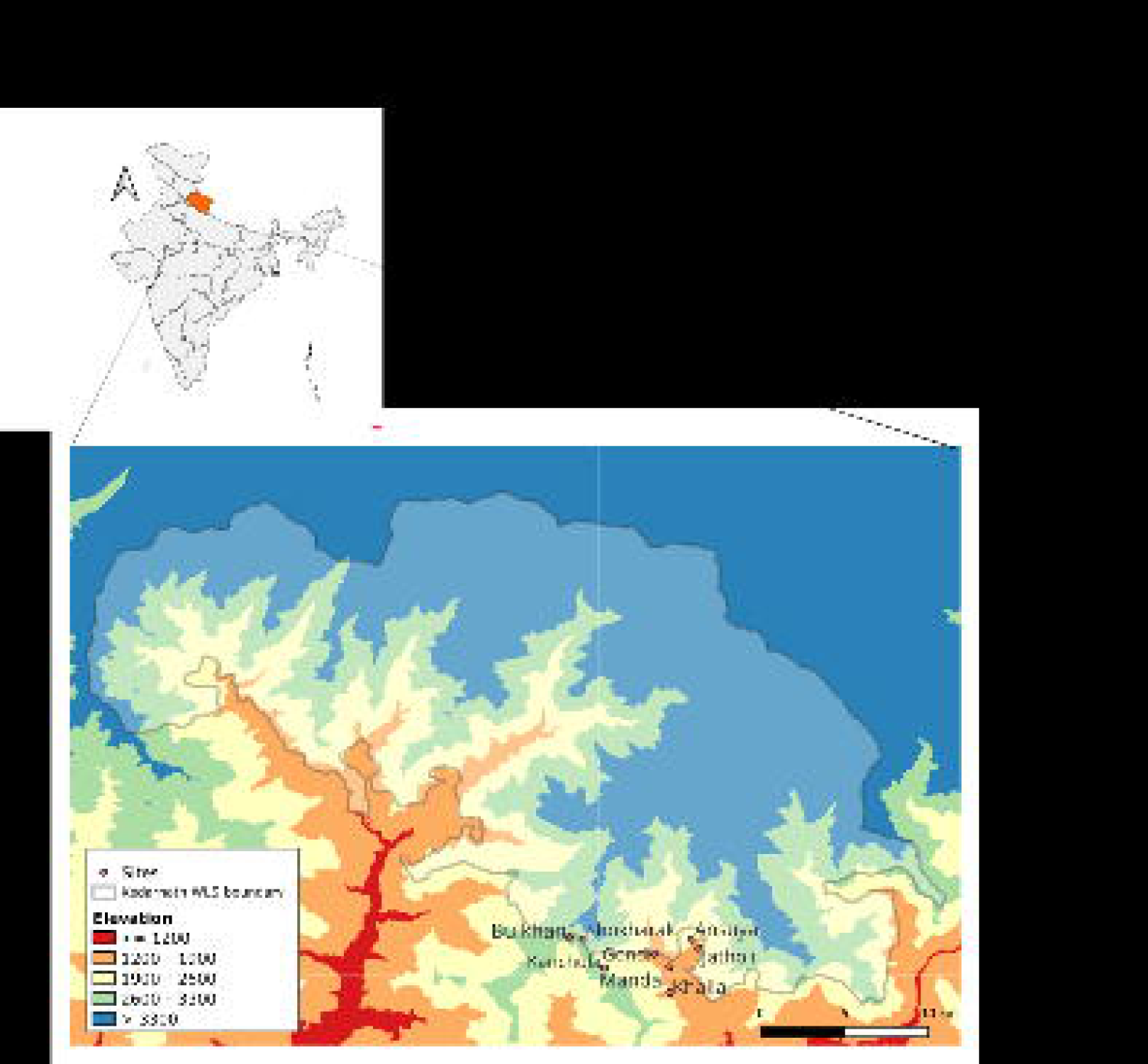
Map showing the sampling locations in and around the buffer zone of Kedarnath Wildlife Sanctuary, which is located in northern side of the mountainous state of Uttarakhand (highlighted on the map of India on the left side)

### Field sampling

The study area was stratified based on elevation and vegetation types to sample the moth diversity along the gradients. Sampling was done every 200m along the elevation from 1500m up to 3100m. For analyses, I categorized the sampling sites based on similar elevation and vegetation types as follows: Khalla, Mandal, Gondi (below 2000m: Low Elevation); Jatholi, Ansuya, Kanchula Kharak (2000-2500m: Mid Elevation); Bulkhan and Shokharak (>2500m: High Elevation). The vegetation types ranged from the Himalayan Moist Temperate Forests dominated by *Quercus* spp.(Low elevation) and mixed Oak Forests (Mid elevation) and; sub-alpine forests with *Rhododendron* spp-Alpine meadows with (High elevation) (Champion and Seth, 1968). The low, medium, and high elevations were therefore used as factors in statistical analyses. Moreover, spacing elevation bands out in this manner also reduces the spatial autocorrelation across our sampling sites.

At each site, 3-night sampling was done (details provided in Table S1 and the sampling sites are shown in Map 1). On each sampling date, two LepiLED (Brehm, 2017) light traps were operated for 3-4 hours after dusk, depending on the weather conditions (Estimated trap hours-Trap 1: 1900-2200; Trap 2: 2000-2300 hrs) which is the peak time of activity of Geometrid moths (Brehm & Fiedler, 2003; and also from personal observations during reconnaissance survey). Also, logistic constraints do not permit all-night sampling. Sampling was done for a total ∼140 hours from March to May (Spring-Summer season) in 2018, during which the average temperatures during sampling were, 16.2 °C, 11°C and 7.5°C at the low, mid and high elevation zones respectively.

The minimum distance between the two light traps was 50 m, with lamps not in line of sight of neighbouring sites, so that cross-site sampling does not occur. Geometrid moths attracted to traps were counted, and representative samples from each species were collected for morphometric measurements and as vouchers. While many species were represented by only one specimen (singletons), an average of three specimens were collected whenever possible for more reliable morphological analysis. All species were included for morphometric measurements, analyses, and diversity assessment. The specimens were collected only from March to May in 2018, as per the permit from the Forest Department. The specimens were then identified till species, as much as possible. The voucher specimens have been submitted to the Research Collections Facility at the National Centre for Biological Sciences, Bangalore, India.

### Morphological trait measurement

All voucher specimens were curated and photographed. I measured morphological traits on the scaled photographs of 697 specimens for morphometry using the software Image J (Schneider, Rasband & Eliceiri, 2012). As different linear measurements widely used proxies for body size in Geometrid moths, (Brehm *et al.,* 2019; Foerster *et al., 2*024)). I measured the body length, wingspan (the width between the apices of the forewings, with the trailing wing edge perpendicular the body), forewing length (measured from the body to the apex of the forewing) and hindwing length. Further, forewing area, hindwing area, were used to calculate Wing loading [body length/ 2*(forewing area R+ hindwing area R)] with body length as a proxy for body mass and total wing area is twice the area of right forewing and hindwing combined (Shi *et al*., 2015), and the ratio of hindwing area to forewing area which is roughly termed manoeuvrability (Arrizabalaga□Escudero *et al,* 2019), (Figure S2) which is directly related to the flight capability of moths. These measured traits are thus predicted to govern the occurrence and sustenance of geometrid moths across elevational gradients.

### Analyses

#### Species diversity

The elevation bands for the sites selected for sampling in my study area were separated by roughly 200 m each. In order to perform meaningful statistics using elevation as a continuous variable, the bare minimum would have been three sites at every 200 m. This was, unfortunately, not possible given the topography of the landscape. Therefore, I pooled the sampling sites into low (1500-1600m; Mandal, Khalla, Gondi), medium (1700-2000m), and high (2600-3000 m). The low, medium, and high elevations were therefore used as factors. I used the ‘iNEXT’ R package (Hsieh *et al.,* 2016) to calculate rarefied species diversity extrapolated to the number of individuals across different elevation classes. This approach allows to visualize species diversity as a function of sampling effort or the number of individuals sampled. By rarefying the data, the species diversity estimates can be standardized across elevation classes, enabling a comparison of species richness among sites with different sampling intensities. To show sample completeness, I similarly calculated the rarefied sample coverage at each of the elevation classes.

To understand the effect of elevation on the species composition in the assemblages, I calculated the overall beta-diversity (Sørensen pairwise dissimilarity), and species spatial turnover (Simpson’s pairwise dissimilarity), all measured as average dissimilarity of the observed units to their group centroid (Anderson *et al*., 2006). This analysis was done with the ‘betapart’ package in R (Baselga & Orme, 2012), which decomposes the total beta diversity into two components, Nestedness (βnested): this component represents the variation in community composition due to species loss or gain across sites that are nested within others. It reflects the turnover of species due to differences in site richness; Turnover (βturnover): this component represents the variation in community composition due to species replacement between sites. It reflects the substitution of species among sites.

#### Variation in the individual trait values, trait dispersion and trait space overlap (niche differentiation)

For understanding the effect of elevation on individual traits, I used two approaches. First, I investigated the response using community abundance-weighted mean trait value (CWM; Lavorel *et al.,* 2008), a widely used method to calculate community trait values weighted by abundance of species in that community. The change in CWM with elevation was quantified using ordinary least squares regression, where each trait was used as a predictor variable in a separate model with Elevation (factor) as the response variable. Then the raw mean of the individual trait values were compared by pairwise comparisons. It is useful to look at both the approaches as the relationships between community-weighted mean (CWM) traits and elevation, is supposed to reflect selection towards locally optimal traits (Violle & Jiang, 2009), but it is limited by the large amount of interspecific trait variation typically found within ecological communities (Muscarella & Uriarte, 2016).

To understand the trait dispersion at the subfamily-level and at the species assemblage level across the elevational gradient, I compared the homogeneity of dispersion (Anderson *et al.,* 2006) for the assemblages in each elevation category. I used the *betadisper* function of the R package vegan (Oksanen *et al*. 2009) to calculate dispersion, which tests whether the variance or dispersion of two or more groups is significantly different or not based on Euclidean distance. This was followed by a *permutest* for the homogeneity of multivariate dispersion. Additionally, I have used Principal Coordinates Analysis (PCoA), to visualize pairwise distances in a two-dimensional plot followed by PERMANOVA.

To understand the overall trait overlap (niche differentiation) among species assemblages across the elevational gradient, I used O-statistic (Read *et al.,* 2018) to generate a single community-level (pooling the assemblages across the elevation) overlap index across multiple traits by fitting non-parametric kernel density functions to the distribution of trait values within an elevation (Mouillot *et al.,* 2005; Geange *et al.,* 2011; Read *et al.,* 2018). For this, body length, wing loading, and Manoeuvrability were chosen as the least correlated independent variable, to avoid multicollinearity. For this, The median pairwise overlap was calculated by first determining the overlap of traits in each species pair, and then taking the median overlap of each species pair in a community. This median overlap value is called the O-statistic (O for overlap)

The R package *Ostats* (Read *et al.,* 2022) was used to calculate O-statistic, the multivariate O-statistic estimation in this package relies on functions imported from the package *hypervolume* (Blonder *et al.,* 2018). To test whether the degree of niche overlap among species in each elevation category is greater than or less than expected by chance, I evaluated the O-statistic against a null model, where the median of the pairwise hypervolume overlaps are weighted against the abundances of the species pairs in each community or Standardized Effect Size (z□scores).

## Acknowledgements

This work would not have been possible without funding from the Rufford Foundation, UK. It was an independent endeavour, and I am immensely thankful to the Dean, Wildlife Institute of India, Dr. V.P. Uniyal and the Uttarakhand Forest Department (Letter no.2921/5-6(WII) dated 17 April 2018) for permitting me to carry out this work on my own. I would like to thank Prabhat Singh Bisht, my local field assistant, and all the volunteers who helped me during the fieldwork, namely: Meghavi Purohit, Kunjan Joshi, Bhargav Bhadresvara, and Mohit. I extend my sincere thanks to Anza Simon and Rama Narayanan for helping with the morphometry of the specimens, which were pinned and curated by Tarun Karmakar. Lastly and most importantly, I am immensely thankful to Saskya Van Nouhuys, Mayank Kohli, and Rohit Chakravarty for their valuable time, support, and discussion regarding the analysis and writing of the manuscript. The manuscript was written and the data was analyzed while I was funded by Dr. D.S. Kothari postdoctoral fellowship (2020-2023), though this fellowship was not used in conducting the research for this paper.

## Data availability statement

The data that support the findings of this study are available from the corresponding author upon reasonable request. Upon the acceptance of the manuscript, the data will be made accessible in a designated data depository.

